# Topological Investigation of Protein Folding and Intrinsic Disorder

**DOI:** 10.64898/2026.02.08.704645

**Authors:** Muriel Elizabeth Hammond, Vasily Akulov, John van Noort, Laura Zwep, Alireza Mashaghi

**Author notes:** co-first authors (Muriel Elizabeth Hammond, Vasily Akulov).

## Abstract

Mapping protein conformations into a space of fold topologies offers an unprecedented perspective on the long-standing protein folding problem. In this study, we apply circuit topology to investigate the folding landscape of both stably folded and intrinsically disordered proteins. This topological approach quantifies intra-chain contact arrangements within a polypeptide chain. We demonstrate that ordered and disordered proteins can be distinguished by their topological organization, and that a topology-based model can predict chain compaction and folding state. Furthermore, topology relates to folding and unfolding kinetics and thermodynamics. These findings establish topology as a fundamental concept for understanding protein folding and disorder.

## 1 Introduction

Protein folding is a fundamental biological process, directly linked to biological function and disease.^1^ Folding of a protein chain leads to an ordered 3D structure that is intricately linked to function; even minor deviations in structure can cause pathology. In contrast, intrinsically disordered proteins (IDPs) or proteins containing intrinsically disordered regions (IDRs) typically lack a stable three-dimensional structure under physiological conditions.^2,3^ Increasing evidence shows that IDPs and fully folded proteins represent opposite ends of a continuous spectrum of structural flexibility.^4^ An important question is how one can describe and quantify this spectrum in a unified framework.

For a long time, it was believed that only well-folded proteins could perform biological function, but emerging data expanded this view. Despite the lack of a single, stable structure, IDPs play crucial roles in many cellular processes. This functional versatility is partly due to their ability to interact with different binding partners. ^5^ IDPs may do so either by undergoing disordered-to-order transitions upon binding or forming a “fuzzy” complex where structural disorder is maintained during the interaction. ^6,7,8^ IDPs play an important role in many biological pathways and disease mechanisms, such as cancer, neuromuscular diseases and amyloidosis, underscoring the importance of understanding their behavior.^2,9,10,11^ Unlike folded proteins, which adopt a well-defined and compact tertiary conformation, disordered proteins exist as dynamic conformational ensembles.^12^ In disordered proteins, chain conformation fluctuates, visiting various metastable states, separated by relatively low free energy barriers. As a result, the function of disordered proteins is less dependent on structure.^2,13^

Traditional structure analysis approaches rely on well-defined folded structures and are therefore inadequate for describing the behavior of disordered proteins.^13,14^ Many experimental and computational tools developed for folded proteins are limited in their ability to analyze IDPs due to their conformational heterogeneity and lack of a single native state.^15,16,17^ The inability to effectively study disordered proteins has led to a knowledge gap regarding their position within the broader topological landscape of the proteome, and how their lack of stable structure relates to sequence, function, and contact organization. A framework that can generically characterize both ordered and disordered proteins within a unified structural context is an essential step toward a comprehensive structure-function paradigm.

A logical approach to address this challenge is to characterize protein conformation using methods that are less sensitive to disorder yet capture key details of the transient fold. Such a “topological” approach seeks shape properties that remain invariant. In disordered proteins, local residue positions fluctuate on hundreds of nanosecond timescales. This dynamic behavior creates a noisy system where detailed coordinate information rapidly changes. Transient intra-chain contacts are key drivers of folding and can persist for hundreds of nanoseconds to microseconds.^18,19,20^ Every contact within a protein structure restricts its fluctuations by constraining the chain into a specific topology, even though the precise structure of the corresponding chain segments may fluctuate widely. This distinction is essential: conformations change rapidly, but their underlying topology changes only when stable, topology-defining contacts are formed or broken. Because of this stability, circuit topology provides a way to follow the dynamics of highly flexible or disordered proteins—not by tracking noisy atomic positions, but by tracking how the set of contact relationships evolves over time. By classifying pairs of contacts as series, parallel, or cross regardless of their sequence separation, circuit topology captures transient structural organization even in systems dominated by disorder. Recently, this framework was used to track the dynamics of a disordered fragment of the glucocorticoid receptor, where an increase in compaction was accompanied by an increase in topological complexity.^21^

Circuit topology (CT) characterizes the arrangement of these intra-chain contacts and may therefore be more robust for conformational analysis of (partially) disordered proteins.^18,19,20^ Focusing on contacts and their arrangements offers a way to reduce this complexity while capturing essential aspects of folding. These relatively stable contact patterns reveal invariant topological features that persist despite rapid chain deformation. By disregarding fast fluctuations, this approach captures the structural essence of both folded and disordered proteins.

CT is a fundamental property of a linear polymer. By representing proteins as interconnected circuits, it captures how intra-chain interactions are arranged.^19,22^ In its simplest form, CT classifies pairs of contacts into three groups: series (S), parallel (P), and cross (X) (Figure 1A). In a series arrangement, contacts are not entangled and do not “interact” from a topological perspective. In a parallel arrangement, one contact is nested within another. In a cross arrangement, the contacts are partially interwoven with each other. Hereafter, these topological contact arrangements are referred to as topological parameters. This mathematical framework can be used to describe how a polymer chain folds, unfolds and rearranges during conformational transitions ^18,20,23,24,25,26^. CT has, for example, already been shown to be capable of predicting the folding rate of proteins.^27^

**Fig. 1.**
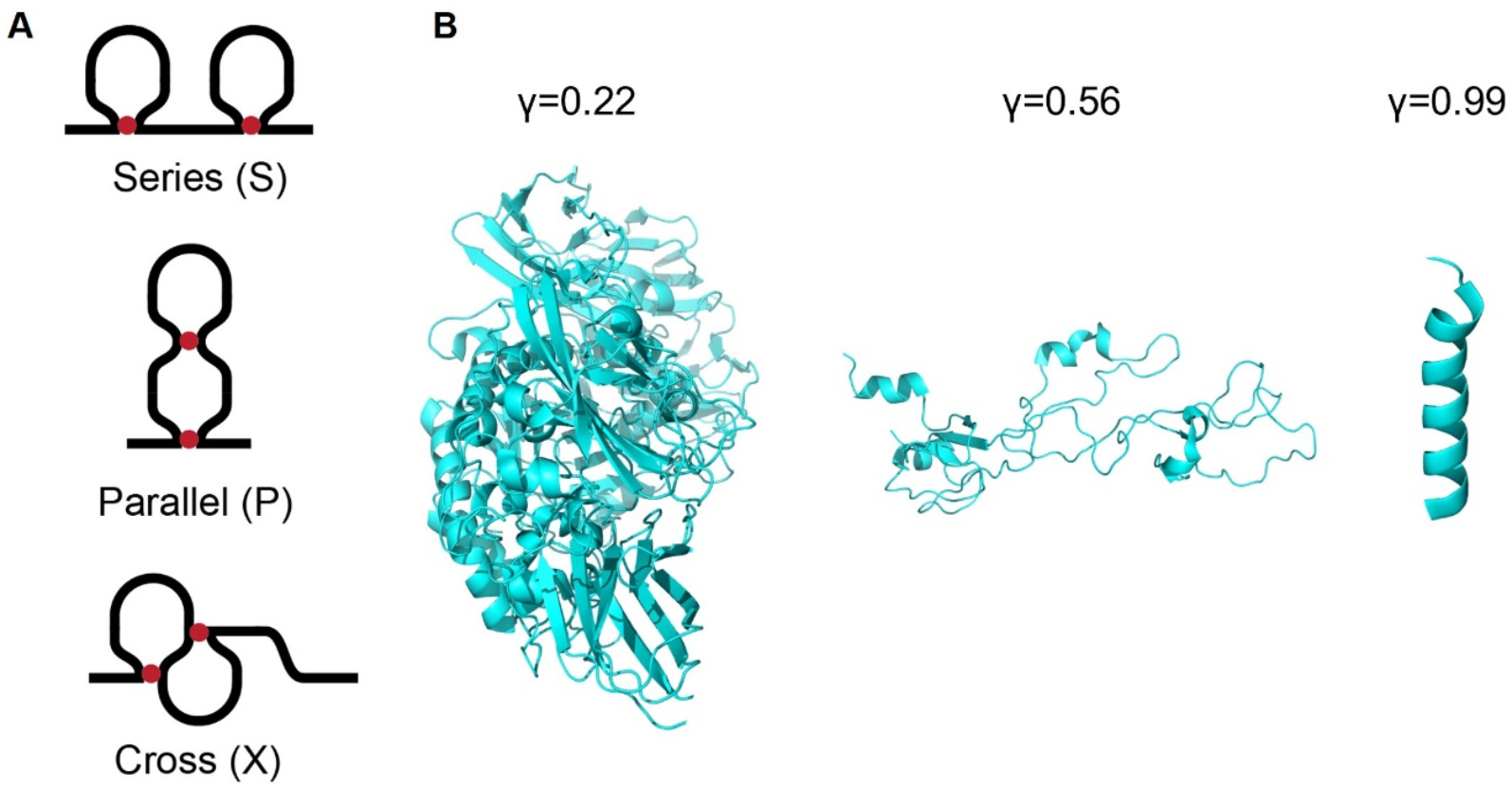
Protein circuit topology and compaction measures. A. The three basic topological arrangements in the circuit topology theory. From top to bottom: series (S), parallel (P), and cross (X) B. γ obtained from the Flory random coil model is used as a folding exponent to characterize the compaction of proteins in the database. A folded protein has a Flory exponent around 0.33 and a more disordered protein around 0.6. Long helical structures often have a γ around 1, although they are folded structures. Proteins from left to right have PDB IDs: 3OG2, 1OCY, 1BTR. The first protein is obtained from the SCOPe database, and the last two are obtained from the PED database. Notably, folded globular (3OG2) and long helical (1BTR) proteins exhibit very different Flory exponents, highlighting the need for a different metric to characterize folding.

Recent research on a small set of IDPs has shown that CT is also valuable for studying proteins whose transient structures are challenging to characterize with conventional techniques.^28,29^ However, a systematic analysis of the universe of identified folded and disordered proteins is still lacking. Visualizing the topological landscape of proteins — e.g., the distribution of proteins along the P, S, and X axes—would provide a new perspective on difficult-to-characterize processes, such as protein folding and disorder-to-order transitions.^7,30^

Here, we take a systematic approach and analyze the topological organization of structured and intrinsically disordered proteins in a unified framework. Topological signature markers of these protein classes are revealed by quantifying differences in their topological composition. These differences are then examined in relation to protein compaction, disorder, folding thermodynamics, and folding kinetics. This analysis delineates the organization of proteins within topological space and clarifies how contact arrangement correlates with folding behavior. Finally, topology-based models are developed to predict and characterize the protein folding landscape across folded and disordered proteins.

## 2 Methods

### 2.1 Computational environment

All computational analyses were performed in Python^31^ (version 3.12.7). The primary Python libraries used included NumPy^32^, Pandas^33^ for data manipulation, Scikit-learn^34^ for machine learning, MDTraj^35^ for structural analysis of molecules, and Plotly^36^ for data visualization.

### 2.2 Databases and data preprocessing

To perform CT analysis of disordered and folded proteins, different protein databases were used to address specific sub-questions, as no single database contained all the required data or sufficient sample sizes for the research. To predict the folding type of proteins, data from the Structural Classification of Proteins - extended (SCOPe 2.08, filtered by *fold*)^37^ and the Protein Ensemble (PED)^38^ databases were obtained. The SCOPe database contains mostly folded proteins, and one of every fold type of the proteins was used for analysis (n=311). The PED database contains mostly disordered proteins and has 219 proteins that could be used for analyses after preprocessing. SCOPe and PED do not provide information on the folding and unfolding rate of the proteins. Therefore, the K-Pro (n=51)^39^, ACPro (n=118)^40^, and ProThermDB databases (n=58).^41^ were used to examine the relationship between the thermodynamics, kinetics, and CT data of proteins. The K-Pro data base contains information on the kinetics of folding and unfolding. ACPro contains additional data on folding kinetics and ProThermDB contains a database of folding free energies. As such, the data from the databases were combined to enable more comprehensive analysis of kinetics and thermodynamics. To train each model, the datasets were divided into training and testing subsets. The exact ratio for each model is provided below. Entries were selected randomly.

After obtaining Protein Data Bank (PDB) files from the databases, a consistent pre-processing method was used for all databases. The PDB files were saved as single structures and filtered to keep single chains. For the thermodynamic analysis, the structures were filtered to retain only protein fragments for which experimental kinetic information is available. Only proteins represented by fewer than 200 structures were used. Protons were removed from the coordinate files. After pre-processing, the circuit topology parameters^18^ and contact order^42^ were calculated for each conformation, and then the averaged values were used for each protein.

### 2.3 Topological analysis

For topological analysis of the proteins, we employed ProteinCT, an open-source Python implementation of the protein circuit topology framework developed by Duane Moes et al. was used.^43^ This enabled single chain circuit topology analysis which categorizes pairs of intra-chain contacts into three categories: parallel (P), series (S) and cross (X). This script takes the PDB files as input and performs CT analysis to obtain the number of arrangements of each category in the protein conformations. Contacts between neighbouring residues are excluded by setting the *exclude neighbours* parameter to two, ensuring a minimum sequence distance of three between contact sites. Other parameters, such as Flory scaling (*γ*), solvent accessible surface area (SASA), and radius of gyration (*R*_*g*_), were obtained using a modified version of a previously published Python script (See Supplementary GitHub repository).^29^

### 2.5 Contact order

The contact order (CO) is a structural property of a protein fold. ^42,44^ The contact order is defined as the average sequence separation of residues that form contacts with each other in the three-dimensional structure.^45^

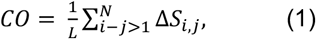

*L* is the total number of residues of the protein. *N* is the total number of contacts and Δ*S*_*i,j*_ is a sequence separation between residues *i* and *j*. ^42^

Proteins with a low contact order tend to form contacts with nearby residues in the sequence, while proteins with a high contact order tend to have interactions that are further spread out over the protein sequence.

### 2.4 Flory random coil model

To define the compaction of a protein, the Flory exponent (*γ*) was used, following the framework defined by Alston et al. In this framework, a protein is seen as a freely jointed polymer chain with excluded volume. The only constraint the polymer has is the distance between monomers.^46^ Following the approach suggested by Rahul K. Das *et al*.,^47^ *γ* was obtained by fitting Equation 2 to the distances between residues within a protein:

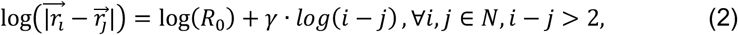

in which 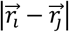 is the distance between residues and *i, j* are the indexes of monomers for all pairs.^46^ Thus, the logarithm of the distance between residues depends linearly on the logarithm of the sequence separation (*i* - *j*), where the Flory exponent *γ* is the slope and the logarithm of distance between monomers, log(*R*_*0*_), is the intercept. This method was applied to both folded and disordered proteins. If a protein has multiple coordinate files, all conformations were used for the calculations. Thus, a single *γ* value per protein was obtained from fitting the Flory random coil model. Example proteins with the corresponding *γ* values are shown in Figure 1B.

A sigmoidal regression was used to correlate topological parameters and the Flory exponent. The data were split into training (424) and testing sets (106) (80:20 respectively) before training the model. Test data were used to evaluate the model’s performance. The accuracy of the model was characterized by the *R*^2^-score. This score represents the proportion of variance in the dependent variable that is explained by the independent variables.

### 2.6 Protein folding prediction model

Logistic regression was used to classify proteins into folded or disordered based on their topological parameters. To train a logistic regression model that predicts whether a protein is folded or not, a “clean” database containing solely folded or disordered regions of proteins was created. From every protein in the SCOPe and PED databases, the longest folded and longest disordered regions were taken. The ‘folding’ of the region was determined based on the secondary structure. Using the Define Secondary Structure of Proteins (DSSP) algorithm, the secondary structures (helix (H) or extended (E) for folded regions, and random coil (C) for disordered regions) were obtained. Folded and disordered regions with a minimum length of five residues and a maximum of three interruptions of a secondary structure of the opposite folding type were saved and used to train the model. The data were separated into the test and train datasets in a proportion of 20:80, (216 structures for testing and 863 structures for training).

### 2.7 Free energy of folding

From the K-ProACPRO databases, data on the unfolding and folding rates for every protein were obtained. Using this data, the free energy of folding was calculated

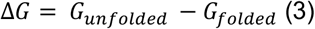

Here, Δ*G* is the free energy difference between the folded and unfolded states of the protein.^42^ The free energy difference relates to the folding and unfolding rates according to:^48^

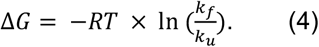

To this free energy prediction were added the data from ProThermDB. From all databases, only natively monomeric proteins that do not contain cofactors (e.g., heme) were selected. Linear regression was used to correlate the free energy of folding, folding rate, and unfolding rate of proteins with protein length, CO, and topological parameters. The data were equally split into training and testing data. The training dataset for the unfolding kinetics contained 26 data points, with 25 data points used for testing. For the folding kinetics model, we used more data, as it combined the K-Pro and ACPro databases, resulting in 59 data points for training and 59 for testing. For free energy of folding prediction model the training dataset contained 53 proteins and 53 for testing. Model accuracy was evaluated using the mean square error (MSE) and the Pearson product–moment correlation coefficient (*R*) calculated with the *SciPy* library. Significance was tested using a two-tailed *p*-value, in all cases it was lower than 0.05.

### 2.8 Regression analysis

All regression analyses were performed using Python and the scikit-learn library.^34^

## 3 Results and Discussion

Structural diversity of both disordered and folded proteins can be described with the language of topology. CT categorizes contact arrangements in a given conformation and assigns one of three types to each pair of contacts: Series (S), Parallel (P), or Cross (X) (Figure 1A). The number of these topological arrangements characterizes protein topology. First, we investigated the relation between CT arrangements and protein compaction and folding. Next, we analysed the predictive power of CT arrangements on protein folding free energy, as well as folding and unfolding rates.

### 3.1 Circuit topology predicts the Flory exponent

Proteins with a higher number of intramolecular contacts generally exhibit greater compaction (Figure 1B).^28^ However, contact number alone provides only a coarse description of protein structure and is insufficient to capture its topological complexity. To achieve a more detailed and informative characterization, we employ CT, which accounts for the specific arrangement and relationships between contacts.

First, we used structures of folded and disordered proteins from the SCOPe and PED databases as input data, and extracted the Flory exponent and circuit topology parameters (Figure 2A). Next, we split the data into training and test sets at an 80–20 ratio and fitted a sigmoidal model on the training data. The input features were the natural logarithm of the topological parameters. In Figure 2A, the predicted versus the calculated Flory exponent, calculated on the test data, is shown. The model explains 59% of the data variation. We did not observe significant deviations in model performance between the training and testing datasets (Table 1). This suggests that there was no overtraining. In Figure 2B, this prediction model is shown in a 3D plot, where the colored field illustrates the predicted outcomes and the distribution of proteins with different compactions can be seen. Using this model, the Flory exponent *γ* can be calculated using Equation 5:

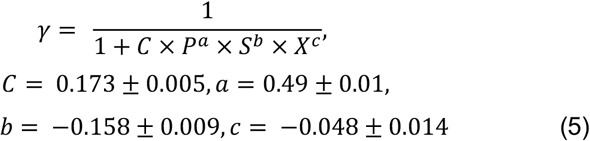

where *P* is the number of parallel arrangements, *S* is the number of series arrangements, and *X* is the number of cross arrangements. *C, a, b*, and *c* are the fitting parameters. In the derivation of this equation, we considered the fact that the input features were the natural logarithm of the topological arrangement numbers. The derivation can be found in the supporting materials (S4).

**Table 1.** Quality of the compaction model.

| Dataset | MSE | R<sup>2</sup> |
| --- | --- | --- |
| Train | 0.0173 | 0.591 |
| Test | 0.0168 | 0.593 |

**Fig. 2.**
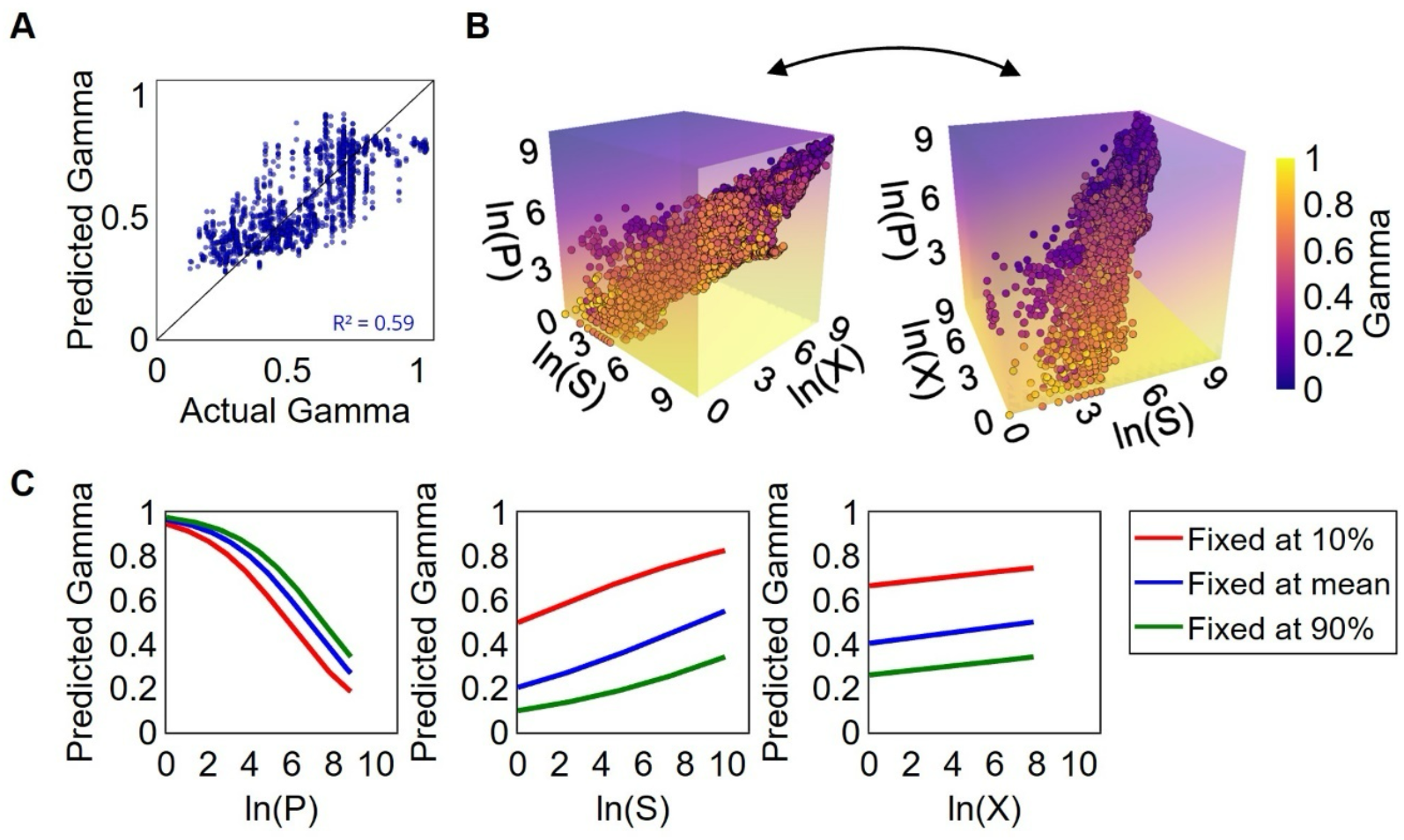
Protein conformational diversity in circuit topology representation. A. Flory exponent correlates with CT parameters with an accuracy of 0.59. In this plot, the predicted γ is plotted against the actual γ. The plot shows that all gamma values are predicted to fall between 0 and 1. Higher γ values are more frequently underpredicted by the model, while a γ values between 0.2 and 0.4 are more often overpredicted. B. Compaction predictions of the PED and SCOPe proteins. This plot shows the distribution of proteins in CT parameter space. The colored field indicates the predicted protein compaction in CT parameter space. Proteins with lower P and higher S and X values are predicted to have higher γ. Disordered proteins have fewer contact arrangements, placing them in the lower part of the plot, while folded proteins have more contacts and are located in the upper part. This demonstrates that the balance between different types of topological arrangements can explain protein compaction. C. Topological parameters show different correlations with the predicted Flory exponent. For each topological arrangement the other two contacts were fixed to the mean, 10th, or 90th percentile of the dataset. Parallel has the largest weight in predicting γ and cross shows the smallest impact.

Results show a difference in the influence of various contact arrangement types in predicting the Flory exponent (Figure 2C). To plot the influence of the arrangement types individually, the remaining two arrangements were held at a constant number. This was done at three different values: the 10^th^ percentile, the mean, and the 90^th^ percentile of the values in the dataset, indicated by the different colors. These results indicate that parallel arrangements most significantly influence the Flory exponent prediction, whereas the number of cross contacts shows the smallest correlation with the predicted value of *γ*. In the case of series contact arrangements, fixing the other arrangement types at low values alters the relationship with the predicted *γ*.

Previous research has shown that parallel contact arrangements are often non-local, characterizing the protein compaction. Cross contacts tend to be local contact arrangements, commonly found in helices.^49,50^ Short-range contacts are typically more important in the early stages of protein folding as they can form and break more quickly.^49^ This may explain why parallel and series contacts are more predictive of protein compaction compared to cross contacts.

We found a negative correlation between the number of parallel contacts and the predicted Flory exponent (Figure 2C), in contrast to series contacts. Usually, the larger the protein, the more intrachain contacts it has, and consequently, the more topological arrangements it exhibits. However, if parallel arrangements exceed series arrangements, *γ* decreases, leading to protein compaction. Conversely, if series arrangements exceed parallel arrangements, the protein behaves as disordered, with a higher Flory exponent. This observation is in line with previous research, which showed that when the number of parallel contacts is relatively low, it may indicate destabilization of the protein structure.^49,51^ The low importance of cross arrangements likely originates from their local character.

### 3.2 Circuit topology categorizes proteins based on their disorder content

In the previous section, we characterized protein compaction with the language of CT. Although that model demonstrates good predictive performance in describing protein compaction, it does not account for the full structural diversity of proteins. For example, it overlooks long helical proteins, which typically exhibit a Flory exponent around 1. An additional model was developed to improve the predictions and to capture folding principles from the CT perspective (Equation 6). To train this second model, a dataset consisting solely of folded and disordered protein regions was used. The natural logarithm of the number of topological arrangements was used as input features for a logistic regression model. In Figure 3A a good separation between disordered and folded proteins can be seen. Validation on the test dataset showed an accuracy of 0.84 (Figure 3A). The model achieved high precision for both folded (0.85) and disordered proteins (0.82), and recall scores of 0.90 and 0.74, respectively. These results indicate that the model is capable of correctly identifying folded and disordered proteins.

**Fig. 3.**
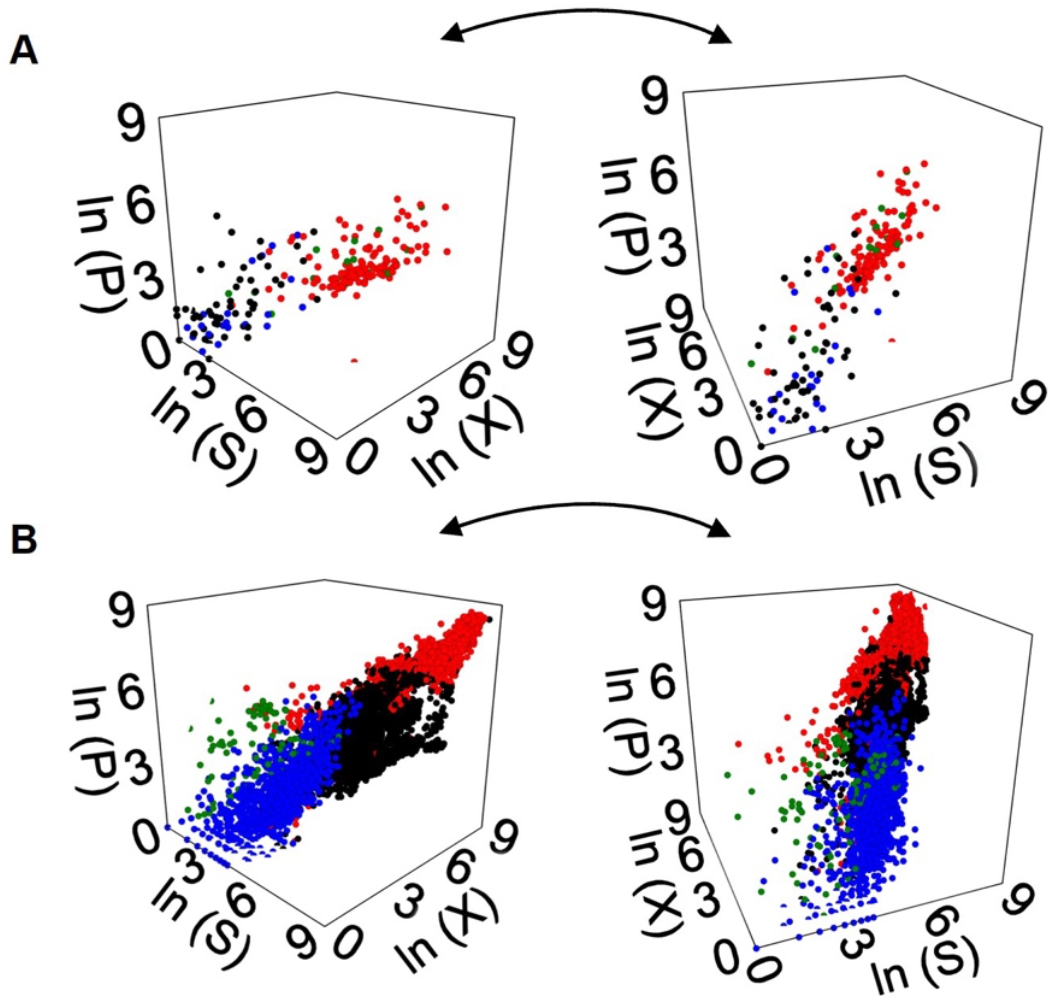
Classification of disordered and folded proteins in circuit topology space. A. The predictions of the test set on the clean database plotted in a 3D plot. Colors indicate whether data points are correctly or incorrectly classified: correctly predicted folded proteins in red, correctly predicted disordered proteins in blue, misclassified disordered proteins (predicted as folded) in black, and misclassified folded proteins (predicted as disordered) in green. B. The predictions of the test set on the PED and SCOPe database plotted in a 3D plot. The same color scheme as in panel A is used. Misclassified folded proteins cluster in one region with γ ≈ 0.95, while proteins misclassified as disordered cluster in another region with γ ≈ 0.75.

Additionally, we validated the performance of the model on the native structures of proteins. We applied this model to predict folding for proteins from the PED and SCOPe databases (Figure 3B). Proteins of these databases were classified to be folded if they had a *γ* lower than 0.4 or higher than 0.9 and disordered if they have a *γ* between 0.6 and 0.9. The model shows clear regions where proteins are correctly classified as folded or disordered, and a distinct area where most misclassifications occur. Proteins that are misclassified as folded, generally have a Flory exponent around 1. Proteins that are misclassified as disordered proteins, generally have a Flory exponent around 0.75 (Figure S1). For this dataset, the model has a predictive power of 54%, with precision for folded proteins at 0.46 and for disordered proteins at 0.91. Recall scores are 0.96 for folded and 0.26 for disordered proteins (Table 2). These lower values likely reflect the continuum of protein structures, as few proteins belong strictly to folded or disordered states. Most proteins contain well-folded regions adjacent to long disordered loops, or adopt multiple conformations that span different degrees of order and disorder. This inherent continuum between folded and disordered states may contribute to the reduced predictive performance, as the model is required to make binary classifications for a biologically continuous property.

**Table 2.** Quality of the classification model.

| Testing data | Class | Precision | Recall |
| --- | --- | --- | --- |
| "Clean" structures | Folded | 0.85 | 0.90 |
|  | Disordered | 0.82 | 0.74 |
| Real protein structures | Folded | 0.46 | 0.96 |
|  | Disordered | 0.91 | 0.26 |

When comparing the models that predict protein compaction and folding, we see a difference in the series and cross contacts parameter. In the “compaction prediction model” (Equation 5), series contacts increase the predicted *γ*, and the effect of cross contact arrangements is close to zero. In contrast, in the “folding classification model” (equation 6) the balance between contact arrangement types is different:

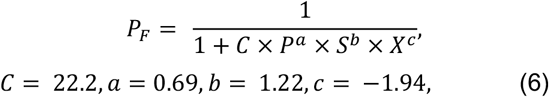

where *P*_F_ – probability for the protein to be folded, P – number of parallel arrangements, S – number of series arrangements, X – number of cross arrangements. *C, a, b*, and *c* are fitting constants, and the input features were the natural logarithm of the topological arrangement numbers. The derivation can be found in Supplementary Materials S4. In this model, parallel and series arrangements decrease the probability for a protein to be folded, whereas cross arrangements increase it. For a protein to be folded, the number of cross arrangements must outperform parallel and series arrangements according to the given coefficients:

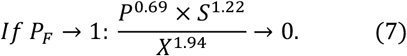

The crucial role of the cross arrangement could be explained by their relatively low count in disordered proteins, while folded structures and long helices typically have more cross contacts per amino acid.^18^

The test set of the clean database shows 84% accuracy in predicting if a protein is folded or disordered. We see the model has a precision score of 0.85 and 0.82 for folded and disordered proteins, respectively. The recall score for folded and disordered protein predictions is 0.90 and 0.74. These results indicate that the model performs very well in both accuracy and completeness for identifying folded versus disordered proteins. By contrast, for the test set of the real protein structure database, the model shows 54% accuracy in predicting if a protein is folded or disordered. We see the model has a precision score of 0.46 and 0.91 for folded and disordered proteins, respectively. The recall score for folded and disordered protein predictions is 0.96 and 0.26.

### 3.3 Circuit topology relates to protein stability

After establishing the ability of CT to quantify protein compaction and folding, we investigated whether topology could give insight into the thermodynamic and kinetic properties of a protein. Equilibrium protein structures are fundamentally driven by minimizing the free energy. The kinetics are defined by the barriers in the free energy landscape describing the folding and unfolding^30^ (Figure 4A). Folded proteins typically display a steep and deep energy well corresponding to a stable native state, while disordered proteins exhibit a flatter energy profile reflecting their conformational heterogeneity.

**Fig. 4.**
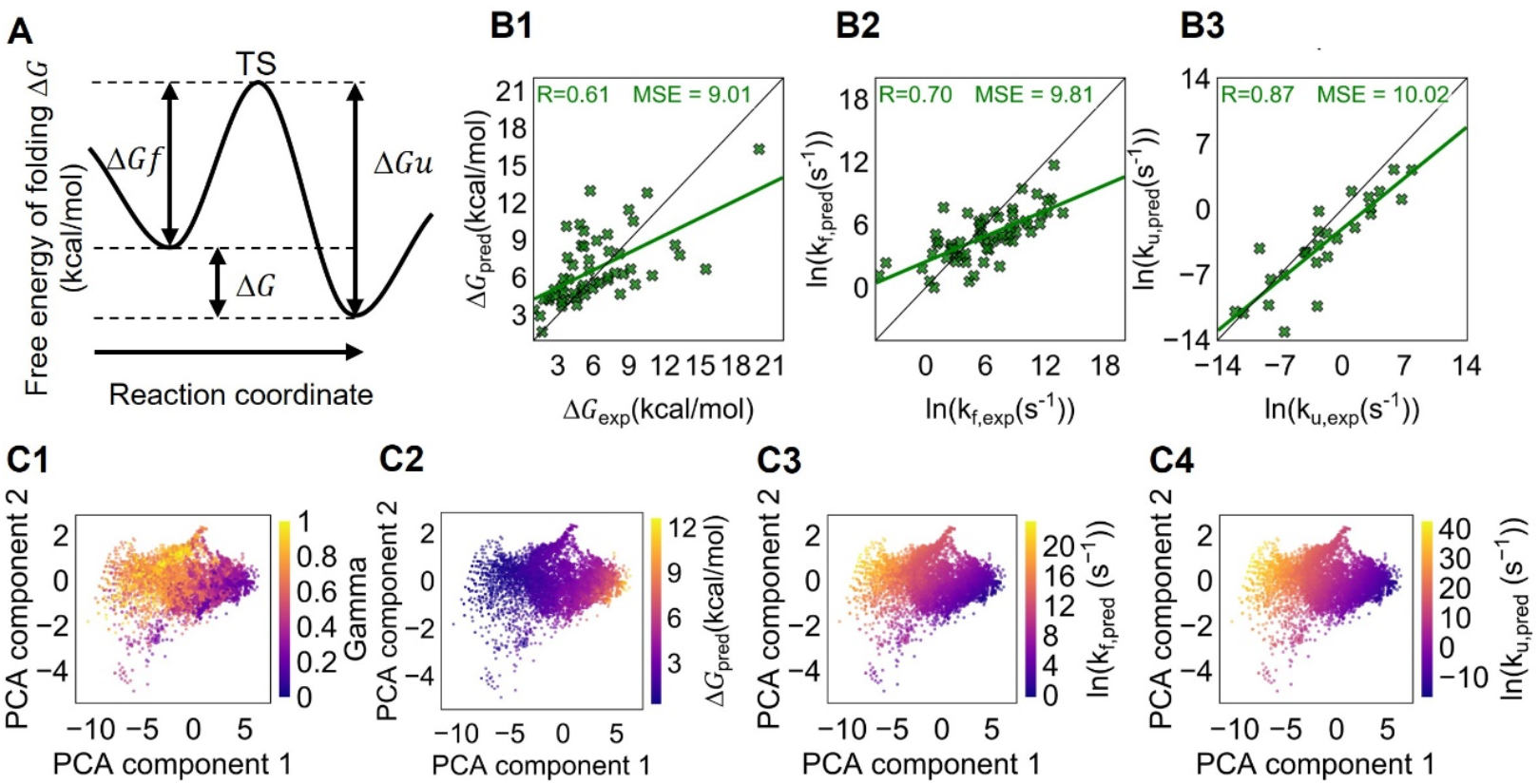
Distribution of protein thermodynamical properties in topological space. A. Visualization of the protein folding profile. The probability of observing a molecule in a particular state is defined by the free energy of that state. The rate of transition between states depends on the height of the energy barrier. B. The model to predict the free energy of folding, unfolding, and folding rate is shown. All plots show the models based on the Circuit Topology (CT) parameters. CT features consistently provide good predictions for folding free energy (B1), as well as for the kinetics of folding (B2) and unfolding (B3). C. The model predicts the protein state and kinetic rates for ordered and disordered proteins. The plots show PCA projections of the CT arrangements for proteins from the SCOPe and PED databases. The plot colored by γ can be used to compare the distribution of folded and disordered proteins with other plots. Folded proteins with lower Flory exponent are more distributed on the right side of the plot, while more disordered proteins are more spread out on the left side of the plot. Plots show, more folded proteins have a higher free energy difference lower folding and unfolding rate compared to disordered proteins.

Since the folding thermodynamics and kinetics are closely linked to how the protein chain is organized in space, we hypothesized that the arrangement of contacts, captured by CT, may carry information on the free energy of folding and folding/unfolding rates. Topological arrangements restrict the polymer chain, with the formation of these restrictions compensating for entropy loss, and different arrangements expectedly affect the chain in distinct ways—this is what this study explores. To explore this possibility, we developed separate predictive models using CT parameters as input features. These models were applied to proteins from the K-Pro and ACPro databases to estimate the free energy difference of folding, as well as the folding and unfolding rates. Prediction models were built using linear regression. Folding and unfolding kinetics, as well as the logarithm of energy, were correlated with the logarithm of topological parameters.

To compare these findings with previous results, the model was also trained with contact order and amino acid length of the protein as input variables. The contact order is a metric that is based on the structure of a protein.^30^ Previous research has shown that the folding rate of small proteins is related to the contact order.^30^ Comparing the results using mean squared error (MSE) and Pearson correlation coefficient for the different models allowed us to evaluate the predictive power of CT features relative to other simpler structural metrics.

This regression model predicts the energy difference between current state (represented by an ensemble of structures) and some completely disordered state. The performance of topological features was compared with protein length and contact order (Figure 4B1). The coefficients, intercept, MSE, Pearson correlation coefficient and, corresponding errors can be found in Table 3. Among these, CT is the only approach that shows good accuracy across all targets—predicting the free energy of folding, folding rate, and unfolding rate.

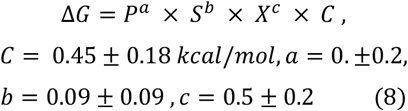

where *ΔG* is the free energy difference between state from the protein structure and some completely disordered state, *P* – number of parallel arrangements, *S* – number of series arrangements, and *X* – number of cross arrangements. *C, a, b*, and *c* are the fitting parameters. In deriving this equation, we accounted for the fact that input features were the natural logarithm of the topological arrangement numbers. The derivation could be found in supplementary material S4. All three arrangement types increase the folding free energy, but parallel contacts have the largest coefficient.

**Table 3.** Model parameters and prediction errors. In this table, all parameters from the prediction models are shown. CT features show better performance than CO for the folding free energy prediction. For kinetics predictions, CT and CO features perform similarly. ΔG is in kcal/mol.

| Predict | Input variables | Coefficients | Intercept | MSE |  | R |  |
| --- | --- | --- | --- | --- | --- | --- | --- |
|  |  |  |  | train | test | train | test |
| $\Delta G$ | Length | 0.024±0.005 | 3.7±0.7 | 9.041 | 14.087 | 0.567 | 0.314 |
| | Contact order | 0.35±0.05 | 1.9±0.8 | 7.346 | 10.825 | 0.670 | 0.519 |
|  | ln(P), ln(S), ln(X) | - | - | 7.495 | 9.012 | 0.683 | 0.605 |
| ln($\Delta G$) | ln(P), ln(S), ln(X) | 0.0±0.2, -0.09±0.09, 0.5±0.2 | -0.8±0.4 | 0.185* | 0.223* | 0.721* | 0.672* |
| ln(kf) | Length | -0.037±0.009 | 8±1 | 9.218 | 13.898 | 0.466 | 0.522 |
| | Contact order | -0.38±0.06 | 9.2±0.8 | 6.622 | 9.456 | 0.661 | 0.744 |
| | ln(P), ln(S), ln(X) | -1.9±1.0, 1.9±0.5, -1.8±1.3 | 16±3 | 6.782 | 9.816 | 0.651 | 0.703 |
| ln(ku) | Length | -0.20±0.03 | 12±3 | 13.670 | 15.812 | 0.762 | 0.720 |
| | Contact order | -1.12±0.16 | 9.6±1.8 | 10.084 | 8.847 | 0.831 | 0.857 |
| | ln(P), ln(S), ln(X) | -4±3, 2.1±1.2, -4±4 | 34±7 | 12.519 | 10.017 | 0.785 | 0.865 |

Results show that CT outperforms CO in predicting the free energy of folding. This highlights the effectiveness of using CT to predict the thermodynamic landscape of protein folding. Despite the dynamic nature of IDPs, the model is capable of predicting the free energy of folding difference. This means the model does not rely on a protein with a well-defined stable structure, but it can capture underlying thermodynamic relationships. In Equation 8 the model is provided as an explicit formula, enabling direct application for estimation.

With equation 8 we predicted the thermodynamic data of the proteins of the SCOPe and PED database (Figure 4C2). This was done to visualize the distribution of free energy across protein topology space. In Figure 4C1, PCA projections of the data points are colored on *γ* to show the distribution of protein folding types. Here, we used the first two PCA eigenvalues, accounting for 92% and 5% of the variability in the data. Comparing 4C1 and 4C2, we observe the co-localization of the compact structures and structures with big free energy of folding. Thus, proteins with a lower *γ* are predicted to have a higher free energy difference relative to the unfolded state (Figure S3C).

### 3.4 Circuit topology relates to protein kinetics of folding and unfolding

To predict the kinetics of folding and unfolding, we built two prediction models, again using the logarithm of topological arrangements as input features. These models were applied to proteins from the K-Pro and ACPro databases to estimate folding and unfolding rates (Figure 4B2 and 4B3):

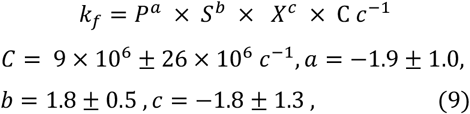

where, *k*_f_ is the folding rate, *P* – number of parallel arrangements, *S* – number of series arrangements, *X* – number of cross arrangements. *C, a, b*, and *c* are the fitting constants. In the derivation of this equation, we considered that input features were the natural logarithm of topological arrangement numbers (see supporting materials S4). Parallel and cross arrangements decrease the folding rate, which is more typical for proteins with complex topology and structure. A similar trend is observed for the unfolding rate:

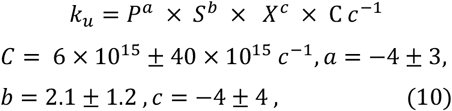

Here, parallel and cross arrangements also slow down the unfolding rate, showing that the same types of arrangements take the most time to form and to break (Figure 4B2 and 4B3).

Next, and for completion, we examined the results of the CT-based models alongside models predicting the folding and unfolding rate based on size (length) and CO, as CO and length are also known to influence folding. The Pearson correlation coefficients of the CT- and CO-based models were comparable (Figure 4B2, 4B3 and supplementary Figure S3A), while the correlation coefficients of the model using length as an input variable was lower (Figure S3B). The MSE results for CT- and CO models were also comparable, while this result was worse for the length model. Consistent results can be obtained for ACPro^52^ database, as studied previously.^49^ This indicates both methods capture kinetic information well, CT via contact arrangement and CO via contact range.^30,49^ However, protein kinetics are limited by the transition state, which remains unknown.^30,53^ One could consider combining these methods to increase the accuracy of predictions.^49^ Another interesting finding was that CT and CO show particularly high accuracy in predicting the unfolding rate, even more so than for the folding rate and free energy of folding. Certain aspects of folding may not be fully captured by the models (Figure 4B2, Figure S3A,B), which can lead to less accurate folding rate predictions for very fast and very slow folding proteins. These results are in line with previous research, which indicated that CT performs somewhat better at predicting unfolding rate than folding rate.^26^ This is explained by a phenomenon called ‘forbidden transitions’, which means that during the unfolding process, a contact cannot be removed because of a parallel contact.^26^ This also reflects differences between folding and unfolding paths: during unfolding, parallel contacts must first be removed, which explains how parallel contacts can slow down the process.^26^ This explains why topology is more effective in predicting the unfolding rate than the folding rate.^26^

We tested the consistency of the folding rate and free energy of folding models. Free energy can be calculated if the folding and unfolding rates are known. Thus, we compared the free energy predictions derived in two ways: direct prediction and recalculation from folding and unfolding rates. As demonstrated in Figure S2, these two methods give very similar predictions, showing the consistency of all models.

We next predicted folding and unfolding rates for proteins in the SCOPe and PED databases (Figure 4C3 and 4C4). In Figure 4C, compact structures cluster with slower predicted folding and unfolding rates. The predicted kinetic parameters also correlate with the Flory exponent (Figure S3D,E). Overall, proteins that adopt compact folded structures are predicted to fold and unfold more slowly than disordered proteins, consistent with the greater topological constraints present in ordered folds.

Topology based models successfully predict both folding and unfolding rates, indicating that the model does not depend on the presence of a well-defined stable structure but instead captures the underlying kinetic principles governing these processes. Equations 9 and 10 present the models in explicit mathematical form, enabling direct application in predictive analysis.

## 4 Conclusions

CT has advanced polymer physics by providing unprecedented approaches to classify and analyze folding, aggregation, structural phase transitions, and entanglement in both single and multichain polymer systems.^20,54,55,56,57,58^ This work establishes CT as a framework for analyzing folded and intrinsically disordered proteins. By quantifying intra-chain contact arrangements, we demonstrate that CT-derived metrics reliably discriminate conformational order, relate to chain compaction, and capture folding thermodynamics and kinetics. This topological perspective extends the utility of polymer physics concepts to protein science, enabling structural and functional inference, even in the absence of stable tertiary structure. Integration of topological approaches with computational modelling and experimental analysis promises to refine models of protein folding landscapes, particularly for disordered states. Furthermore, the delineation of topological signatures in protein ensembles offers a foundation for rational drug design strategies targeting both ordered and disordered protein regions.

Future studies may consider refining the reference model for protein compaction. While the Flory exponent captures general scaling behavior, it simplifies protein structure by assuming random-coil characteristics, with no sequence constraints, other than excluded volume and bond lengths. Combining CT analysis with advanced machine learning and molecular dynamics simulations could yield more detailed and accurate descriptions. Additionally, expanding the dataset of thermodynamic and kinetic parameters would improve statistical power. Incorporating complementary features such as contact order may also enhance predictive performance and provide a more comprehensive understanding of protein folding and disorder. It should be noted that our current approach does not explicitly account for biological or sequence-level factors that can influence protein topology. For example, macromolecular crowding can substantially modulate the structure and dynamics of proteins, as suggested by previous studies on polymer topology under confinement ^55,59,60^ and folding/unfolding of proteins in crowded cytosolic environments.^61^ Similarly, single-point mutations may locally alter topology, and prior work using decision trees and Auto-ML has shown the importance of mutations and post-translational modifications in pathogenicity prediction,^62,63,64^ highlighting areas for future investigation. Finally, our analysis only used a first-order circuit topology approach, while the full circuit topology of a protein includes higher order information (e.g., examining circuit structures^65^) beyond frequencies of pairwise arrangements. Future studies may also apply circuit topology to characterize chain crossings, including knots, slip-knots, and more complex entanglements^66,67,68^. Collectively, our findings provide a topological perspective that bridges structure and dynamics across the protein order–disorder continuum, offering a foundation for future interdisciplinary research in structural biology and molecular design.

## Author contributions

Conceptualization: A.M., and V.A. Project administration and Supervision: A.M. Investigation: M.H., V.A. Formal analysis: M.H. and V.A. Software: M.H., V.A. Methodology: M.H., V.A., L.Z., and A.M. Visualization: M.H. and V.A. Writing & draft preparation: M.H., V.A., and A.M. Review & editing: M.H, V.A., J.v.N., L.Z. and A.M.

## Acknowledgements

The authors thank Jeremy Schmit (University of Kansas) for helpful discussions and acknowledge Leiden University for financial support.

## Supplementary Information

Supplementary figures are provided in the Supplementary Information file.

**Supplementary Figure S1:** Performance metrics for the protein folding prediction model.

**Supplementary Figure S2:** Validation of folding free energy, folding rate, and unfolding rate prediction models.

**Supplementary Figure S3:** Correlation of predicted folding energetics and kinetics with protein length, CO and protein compactness.

**Supplementary Equations:** Equations relating folding kinetics and thermodynamics to topology measures.

## Data availability

Supplementary codes and data associated with this analysis are available at the following GitHub repository: https://github.com/TheMashaghiLab/Publication_Topological_Investigation_of_Protein_Folding_and_Intrinsic_Disorder

## Table of Contents (TOC)

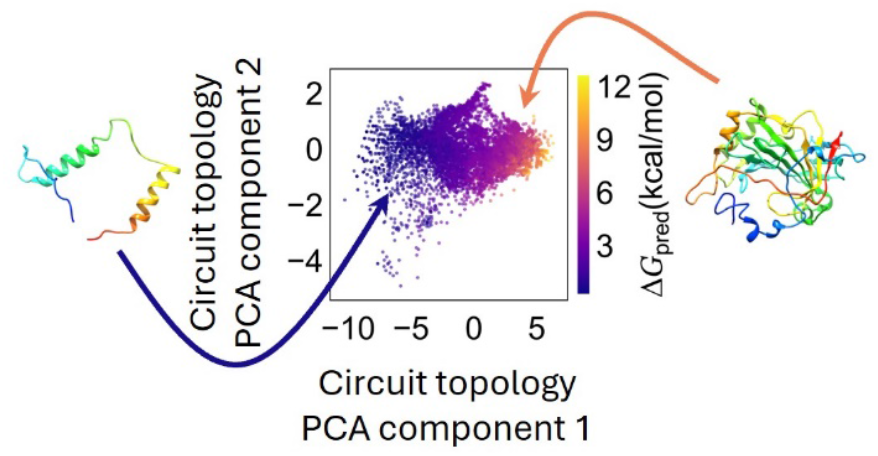

