## Supplementary Information for "Topological Investigation of Protein Folding and Intrinsic Disorder"

[#] co-first authors.

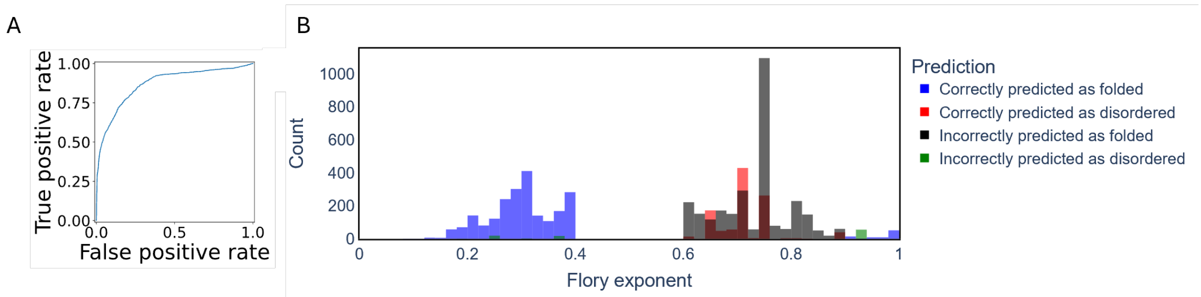


***Supplementary Figure S1*** *Performance metrics for the protein folding prediction model.*

*(A) ROC curve for the protein folding prediction model ("clean" structures dataset), AUC = 0.87.*
*(B) Protein folding model performance on the real protein structures dataset. In this test, the model tends to predict disordered proteins as ordered.*

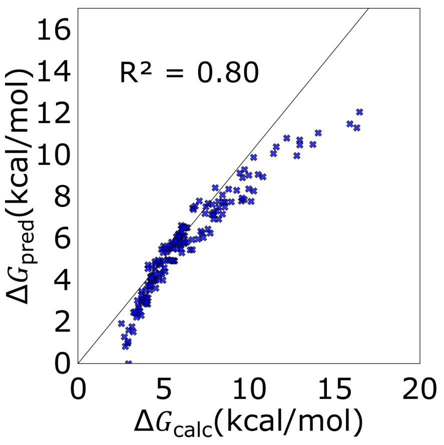


***Supplementary Figure S2:*** *Validation of folding free energy, folding rate, and unfolding rate prediction models consistency.*

*Free energy of folding model scores well on the generalisability test. Model prediction of the free energy of folding based on topological parameters shows comparable results to the calculated predicted free energy of folding. The predicted free energy of folding is calculated using the following formula:* $\Delta G = RT \times ln\ \frac{kf}{ku}$. *The predicted values for the unfolding rate and folding rate from Equation 11 and 12 are used for this calculation.*

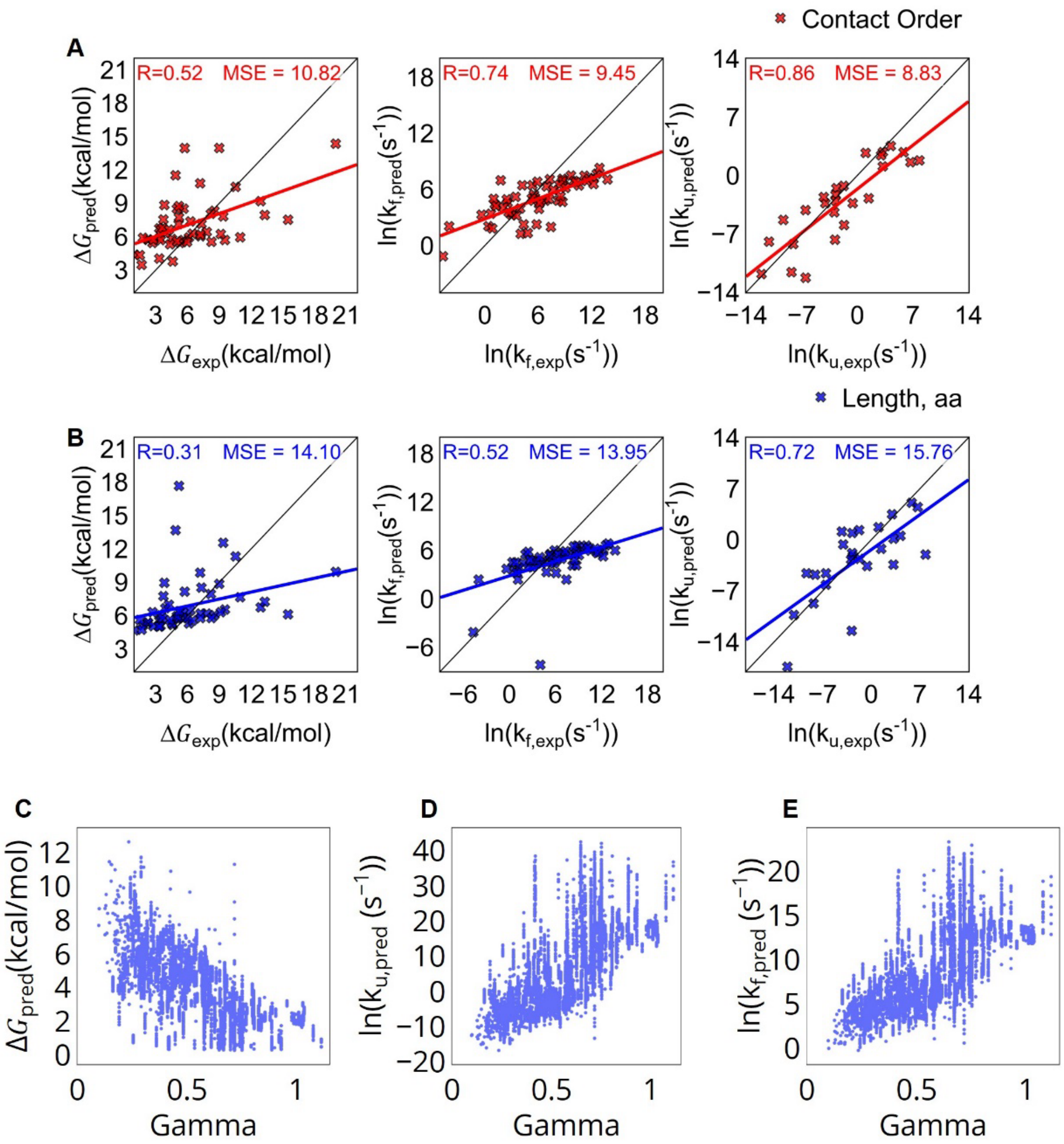


***Supplementary Figure S3:*** *Correlation of predicted folding energetics and kinetics with protein length, CO, and protein compactness.*

*(A) Comparison between predicted and experimental values for a model based on CO. From left to right, free energy of folding, folding rate, and unfolding rate.*
*(B) Comparison between predicted and experimental values for model based on protein length. From left to right, free energy of folding, folding rate, and unfolding rate. In A and B, MSE – mean square error, R - Pearson product–moment correlation coefficient.*
*(C) Proteins with a lower Flory exponent have a higher predicted difference in folding free energy. This means that proteins that are more folded have a higher folding free energy difference from an unfolded state compared to disordered proteins.*
*(D,E) Proteins with a lower Flory exponent have slower predicted folding and unfolding rates than proteins with a higher Flory exponent. This means proteins that are more folded will take longer to fold or unfold compared to more disordered proteins.*

***Supplementary Equations.*** *Equations relating folding kinetics and thermodynamics to topology measures.*

$$\gamma = \frac{1}{1 + e^{\ln(C) + a \cdot \ln(P) + b \cdot \ln(S) + c \cdot \ln(X)}} = \frac{1}{1 + C \times P^a \times S^b \times X^c}$$

$$P_F = \frac{1}{1 + e^{\ln(C) + a \cdot \ln(P) + b \cdot \ln(S) + c \cdot \ln(X)}} = \frac{1}{1 + C \times P^a \times S^b \times X^c}$$

$$\ln(\Delta G) = \ln(C) + a \cdot \ln(P) + b \cdot \ln(S) + c \cdot \ln(X) \rightarrow$$
$$\rightarrow \Delta G = e^{\ln(C) + a \cdot \ln(P) + b \cdot \ln(S) + c \cdot \ln(X)} = P^a \times S^b \times X^c \times C$$

$$\ln(k_f) = \ln(C) + a \cdot \ln(P) + b \cdot \ln(S) + c \cdot \ln(X) \rightarrow$$
$$\rightarrow k_f = e^{\ln(C) + a \cdot \ln(P) + b \cdot \ln(S) + c \cdot \ln(X)} = P^a \times S^b \times X^c \times C$$

$$\ln(k_u) = \ln(C) + a \cdot \ln(P) + b \cdot \ln(S) + c \cdot \ln(X) \rightarrow$$
$$\rightarrow k_u = e^{\ln(C) + a \cdot \ln(P) + b \cdot \ln(S) + c \cdot \ln(X)} = P^a \times S^b \times X^c \times C$$
